# Targeted surveillance strategies for efficient detection of novel antibiotic resistance variants

**DOI:** 10.1101/2020.02.12.946533

**Authors:** Allison L. Hicks, Stephen M. Kissler, Tatum D. Mortimer, Kevin C. Ma, George Taiaroa, Melinda Ashcroft, Deborah A. Williamson, Marc Lipsitch, Yonatan H. Grad

## Abstract

Genotype-based diagnostics for antibiotic resistance represent a promising alternative to empiric therapy, reducing inappropriate and ineffective antibiotic use. However, because such assays infer resistance phenotypes based on the presence or absence of known genetic markers, their utility will wane in response to the emergence of novel resistance. Maintenance of these diagnostics will therefore require surveillance designed to ensure early detection of novel resistance variants, but efficient strategies to do so remain to be defined. Here, we evaluate the efficiency of targeted sampling approaches informed by patient and pathogen characteristics in detecting genetic variants associated with antibiotic resistance or diagnostic escape in *Neisseria gonorrhoeae*, focusing on this pathogen because of its high burden of disease, the imminent threat of treatment resistance, and the use and ongoing development of genotype-based diagnostics. We show that incorporating patient characteristics, such as demographics, geographic regions, or anatomical sites of isolate collection, into sampling approaches is not a reliable strategy for increasing variant detection efficiency. In contrast, sampling approaches informed by pathogen characteristics, such as genomic diversity and genomic background, are significantly more efficient than random sampling in identifying genetic variants associated with antibiotic resistance and diagnostic escape.

## Introduction

Nucleic acid-based diagnostics that enable rapid pathogen identification and prediction of drug susceptibility profiles can improve clinical decision-making, reduce inappropriate antibiotic use, and help address the challenge of antibiotic resistance ^1–3^. However, the sensitivity of such diagnostics may be undermined by undetected genetic variants ^4–12^. Pathogen surveillance programs aimed at early detection of novel variants are crucial to ensuring the clinical utility and sustainability of these diagnostics.

Use of traditional nucleic acid amplification tests (NAATs) for pathogen identification and genotype-based diagnostics for antibiotic resistance can select for genetic variants that escape detection ^13^. Mutations and/or deletions at the NAAT target locus that cause an amplification failure have arisen in *Neisseria gonorrhoeae, Chlamydia trachomatis, Staphylococcus aureus*, and *Plasmodium falciparum*, resulting in false negative diagnostic errors only detected when using another diagnostic platform ^5–7,11^. Diagnostic escape associated with genotype-based diagnostics for antibiotic resistance are the result of resistance-conferring variants (*e.g.*, mutations or accessory genes) not accounted for in the diagnostic’s panel of resistance markers ^4^ and require phenotypic testing to be uncovered.

We recently presented a framework to quantify the sampling rate for early detection of novel antibiotic resistance variants, defining the number of isolates that would need to undergo confirmatory phenotyping from those predicted by genotype to be ^14^ susceptible. Underlying this model are assumptions of unbiased sampling across a population and independence among all isolates. However, these assumptions may not hold in practice, as some subsets of the population (*e.g.*, demographics and/or geographic regions) may be more likely to be sampled than others, and clonal transmission may result in repeated sampling of closely related isolates ^15–18^. The real-world application of this model may also be challenging for pathogens with high case incidence, such as *N. gonorrhoeae*, as the cost of phenotyping required by this model for timely detection of novel resistance variants is likely to be high ^14^.

Implementing a practical surveillance system thus requires improving efficiency over unbiased testing by prioritizing samples in which novel diagnostic escape variants are most likely to be found. There are numerous hypotheses for how to focus sampling and most quickly identify these variants. Novel variants may be more likely to emerge or spread in certain anatomical niches, demographics, or geographic regions ^19–22^, some of which may be systematically under-sampled ^23^ and thus may provide a basis for sampling priority. Data on such characteristics may be obtained from metadata recorded during clinical encounters. Alternatively, they may be inferred from pathogen genomic data. Isolates or clades that are genetically divergent from the majority of isolates in a population may reflect travelers, their contacts, or otherwise under-sampled lineages ^24–27^. Some pathogen genomic backgrounds may be more conducive to the evolution of novel resistance mechanisms ^28^, and markers of these genomic backgrounds (*e.g.*, variants associated with a range of resistance mechanisms and/or resistance to other drugs) may help improve sampling efficiency. Similarly, given historical patterns of antibiotic use, novel resistance may emerge on a background of existing resistance ^29^. Thus, genetic markers of resistance to certain drugs may facilitate identification of lineages more likely to have experienced selective pressures leading to emergence of novel resistance variants.

Here, we test the performance of sampling strategies informed by these hypotheses using *N. gonorrhoeae* surveillance data. *N. gonorrhoeae* offers a useful model, given the increasing drug resistance and recent focus on developing sequence-based resistance diagnostics ^2,30^. We present targeted sampling approaches informed by patient (*i.e.*, demographics, anatomical site of isolate collection, geographical region, recent travel history, or sex worker status) and pathogen (*i.e.*, phylogenetic or genomic background) information. We assess the efficiency of each of these strategies to detect rare (<10% prevalence) resistance variants associated with current or recent first-line recommended antibiotics (*i.e.*, azithromycin [AZM] and extended spectrum cephalosporins [ESCs]), as well as rare genetic variants associated with diagnostic escape, across five genomic surveys with various demographic, geographic, and temporal ranges. We show that phylogeny- and genomic background-aware sampling approaches can increase the detection efficiency of known variants over random sampling, whereas patient feature-based sampling approaches do not. Our results suggest that implementation of such targeted sampling approaches into surveillance programs may reduce the number of cases of novel resistance that occur before it is detected, as well as the resources required to undertake surveillance, compared to random sampling of a population.

## Results

### Composition of the datasets

The datasets (**Table 1**) were biased across patient demographics and/or geographic regions (**Tables S1** and **S2**). Isolates from men and men who have sex with men (MSM) were overrepresented in datasets 1 and 2 compared to overall gonorrhea incidence in men and MSM in the US and Australia, respectively, during the study periods (**Table S2**, *P* < 0.001 for both datasets by chi-squared test of men vs. women and MSM vs. non-MSM in dataset vs. reported incidence). Dataset 4 was comprised exclusively of isolates from men ^31^. While it is difficult to estimate the prevalence of pharyngeal gonococcal infections, as they tend to be asymptomatic ^32^, pharyngeal isolates represented 4% and 18% of isolates with reported anatomical site of collection in datasets 1 and 2, respectively. This suggests either sampling bias across anatomical sites in at least one of the datasets or substantial variation across the two study populations in prevalence of pharyngeal gonococcal infections. Similarly, the geographic distribution of isolates in dataset 3 was significantly different from the reported case incidence across countries (**Table S2**, *P* < 0.001 by chi-squared test of prevalence for each of the countries in dataset 3 vs. the reported overall incidence for each of the countries).

**Table 1.** Summary of datasets.

| Dataset | Temporal range | N<sub>isolates</sub> | Geographic range | Metadata available | SRA study ID/Reference |
| --- | --- | --- | --- | --- | --- |
| 1 | 2011-2015 | 896 | New York, NY, US | Gender, sexual behavior, anatomical site of isolation | ERP011192 [Mortimer et al., 2020, *in preparation*] |
| 2 | 2016-2017 | 2186 | Victoria, Australia | Gender, sexual behavior, anatomical site of isolation, travel history, sex worker status | SRP185594 <sup>33</sup> |
| 3 | 2013 | 1054 | Europe | Country of sample collection | ERP010312 <sup>34</sup> |
| 4 | 2015 | 244 | Japan | Prefecture of sample collection | DRP004052 <sup>31</sup> |
| 5 | 2014-2015 | 398 | New Zealand | N/A | SRP111927 <sup>35</sup> |

### Targeted sampling based on patient characteristics

We investigated whether sampling evenly across demographic groups (demography-aware sampling), anatomical sites of isolate collection (niche-aware sampling), and geographic regions (geography-aware sampling) increased detection efficiency of resistance variants by ameliorating some of the demographic, niche, or geographic sampling biases. We further investigated whether preferentially sampling patients with recent overseas sexual encounters or recent sex work, two characteristics hypothesized to be associated with the introduction and/or increased transmission of resistance ^19,21,22^, increased the detection efficiency of resistance variants. To do so, we simulated and compared the detection efficiency of three genetic resistance variants (**Table 2**) using each of these targeted sampling strategies and random sampling.

**Table 2.** Summary by dataset of the prevalence and distribution of the genetic markers of resistance and resistance phenotypes tested.

| Variant |  | Genetic |  |  | Phenotypic |  |
| --- | --- | --- | --- | --- | --- | --- |
|  |  | RplD G70D | 23S rRNA<br>C2611T (2-4<br>alleles) | *penA* XXXIV | CRO-RS<br>(≥0.12 µg/mL) | CFX-R (>0.25<br>µg/mL) |
| **Drug** |  | AZM<sup>36</sup> | AZM<sup>37</sup> | ESCs<sup>38</sup> | N/A | N/A |
| **Prevalence**<br>**of variant in**<br>**dataset** | **1** | 10.04%<sup>a</sup> | 0.11% | 5.25% | 1.47% | 0.11% |
|  | **2** | 1.14% | 1.24% | 1.69% | 0% | 0% |
|  | **3** | 2.47% | 0.95% | 15.68%<sup>a</sup> | 1.04% | 0.76% |
|  | **4** | 11.07%<sup>a</sup> | 1.23% | 0.41% | 6.56% | 8.20% |
|  | **5** | 0.75% | 0.50% | 2.26% | 0.25% | 0% |
| **Phylogenetic**<br>**D statistic**<br>**for variant in**<br>**dataset** | **1** | -0.18 | 17.50 | -0.29 | N/A | N/A |
|  | **2** | -0.10 | 0.46 | -0.24 | N/A | N/A |
|  | **3** | 0.05 | 0.30 | -0.20 | N/A | N/A |
|  | **4** | -0.16 | 1.83 | 1.81 | N/A | N/A |
|  | **5** | 0.83 | 1.12 | -0.15 | N/A | N/A |

<sup>a</sup>Given the >10% prevalence of RplD G70D in datasets 1 and 4 and *penA* XXXIV in dataset 3, these variants were excluded from sampling simulations.
AZM, azithromycin; ESC, extended-spectrum cephalosporin; CRO-RS, ceftriaxone reduced susceptibility; CFX-R, cefixime resistance

The detection efficiency was not improved by demography-, niche-, geography-aware sampling compared to random sampling for any of the resistance variants (**Table S3, Fig. 1**). In several cases, detection efficiency significantly decreased in demography- or geography-aware sampling compared to random sampling, reflecting enrichment of the resistance variant in the overrepresented demographic or geographic region. However, no significant association between a given resistance variant and demographic group was observed across both dataset 1 and dataset 2, and no demographics or geographic regions were significantly enriched for all variants (**Table S1**), suggesting that preferential sampling of any of these demographics or geographic regions would not be a reliable strategy for increasing novel variant detection efficiency. For example, while *penA* XXXIV was significantly enriched in MSM compared to men who have sex with women and women who have sex with men (MSW/WSM) patients in dataset 2 (*P* < 0.003, Fisher’s exact test), there was no significant difference in the proportions of MSM and MSW/WSM with *penA* XXXIV in dataset 1 (*P* = 0.461, Fisher’s exact test). Similarly, while the AZM-R-associated RplD G70D mutation in dataset 3 was at highest prevalence in patients from Malta and Greece (10% and 6.25%, respectively) and absent from patients from Denmark, the AZM-R-associated 23S C2611T variant was at highest prevalence in patients from Denmark (5.45%) and absent from patients from Malta or Greece.

**Figure 1.**
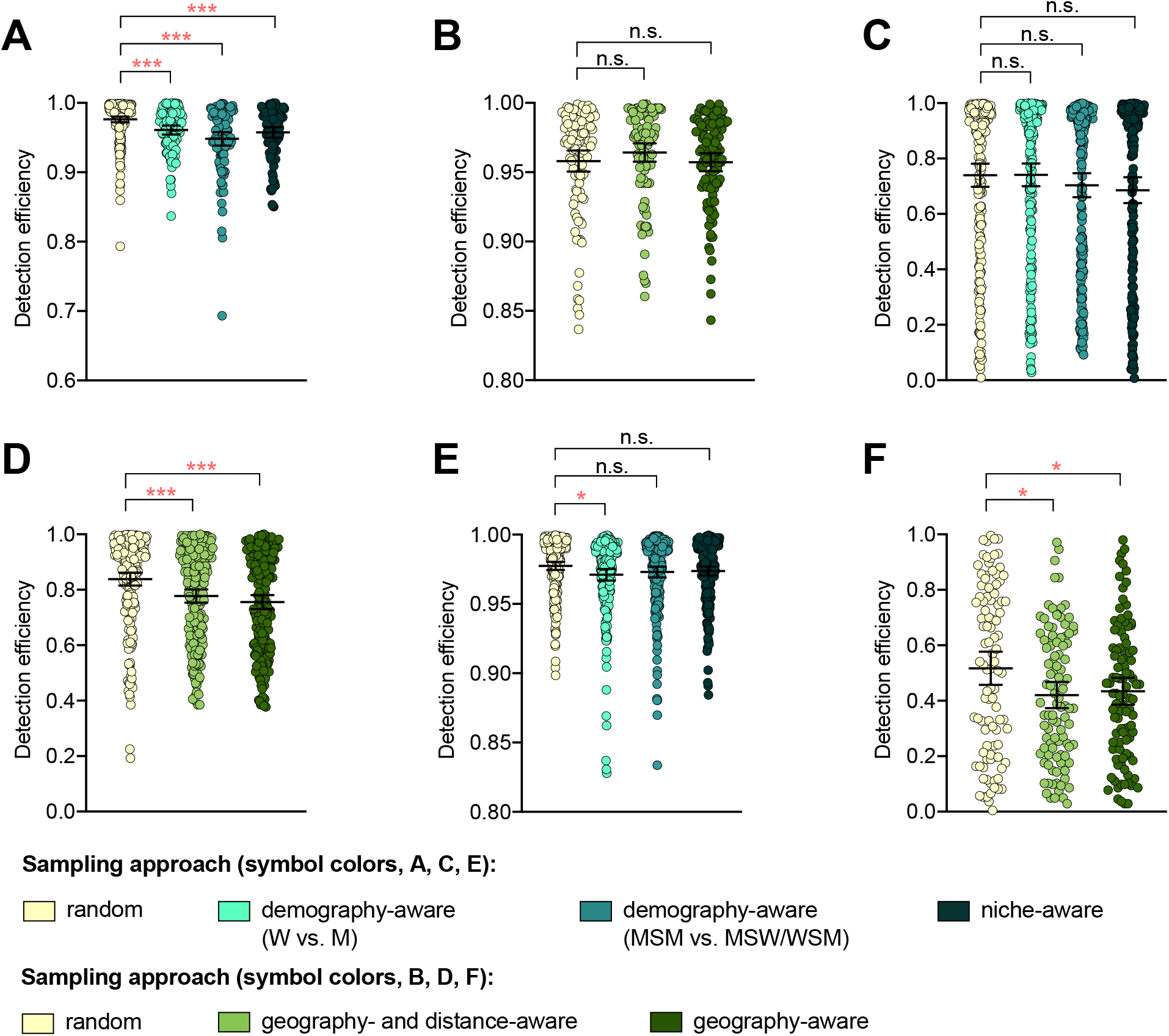
The impact of demography-, niche-, and geography-aware sampling on the detection efficiency of genetic resistance variants. Dot plots showing the detection efficiency (with lines indicating the mean and 95% confidence intervals from 100 simulations) for resistance variants RplD G70D (**A-B**), 23S rRNA C2611T (**C-D**), and *penA* XXXIV (**E-F**) in datasets 1 and 2. In datasets 1 and 2, targeted sampling was informed by demographic (gender and sexual behavior) and anatomical site of isolate collection (niche) information (**A, C,** and **E**), and in datasets 3 and 4, targeted sampling was informed by country or prefecture of sample collection (**B, D,** and **F**). Dot colors indicate the sampling approach, and asterisks indicate a significant difference (*P* < 0.05 by Mann-Whitney U test) in detection efficiency between the demography-, niche- or geography-aware approach compared to random sampling (\**P* < 0.05, \*\**P* < 0.01, \*\*\**P* < 0.001; red asterisks indicate significantly lower detection efficiency of demography- or geography-aware approaches compared to random sampling). Note that sampling simulations were not performed for RplD G70D in datasets 1 and 4 or for *penA* XXXIV in dataset 3 as prevalence of the variants in these datasets was >10%. n.s., not significant at α = 0.05; M, men; W, women; MSM, men who have sex with men; MSW, men who have sex with women; WSM; women who have sex with men.

Isolates from patients with recent overseas sex were associated with significantly longer terminal branches compared to patients that had only engaged in sex locally (**Fig. S1**), in support of the hypothesis that international travel may be associated with the importation of novel or divergent strains, or, more generally, that isolates from travelers may be more likely to be associated with under-sampled lineages. Preferentially sampling from patients with recent overseas sex significantly improved detection efficiency of the RplD G70D mutation and the *penA* XXXIV allele, as these were at marginally higher prevalence in isolates from patients with recent overseas sex compared to those from patients who had only engaged in sex locally (3.03% overseas vs. 0.98% local and 2.02% overseas vs. 1.67% local, respectively, *P* = 0.090 and 0.683, respectively, by Fisher’s exact test for both variants). In contrast, the 23S C2611T mutation was exclusively present in isolates from patients who had engaged in sex locally (**Tables S1** and **S4**). Similarly, while the 23S C2611T mutation was marginally enriched in isolates from patients who had engaged in recent sex work compared to patients who had not (2.33% in sex workers vs. 1.31% in non-sex workers, *P* = 0.327 by Fisher’s exact test), and thus preferentially sampling from sex workers significantly improved detection efficiency of this variant compared to sampling from the full patient population, detection efficiencies for the RplD G70D mutation and the *penA* XXXIV allele were not significantly improved by preferentially sampling from sex workers (**Tables S1** and **S4**).

Together, these results suggest that while targeted sampling based on patient characteristics may increase detection efficiency of some novel variants, it is difficult to predict which groups to target for all potential novel variants.

### Targeted sampling based on genetic diversity

To assess whether preferential sampling of lineages that are divergent from those that have been previously sampled may increase detection efficiency of genetic resistance variants over random sampling, we simulated phylogeny-aware sampling using two methods: 1) maximization of the phylogenetic distance covered with each isolate sampled (distance maximization) and 2) even sampling across phylogenetic lineages (clonal group).

While the distance maximization approach increased detection efficiency compared to random sampling for some variants, there were numerous instances in which this approach, which led to preferential sampling of isolates associated with long branches, substantially decreased detection efficiency (**Fig. 2, Table S5**).

**Figure 2.**
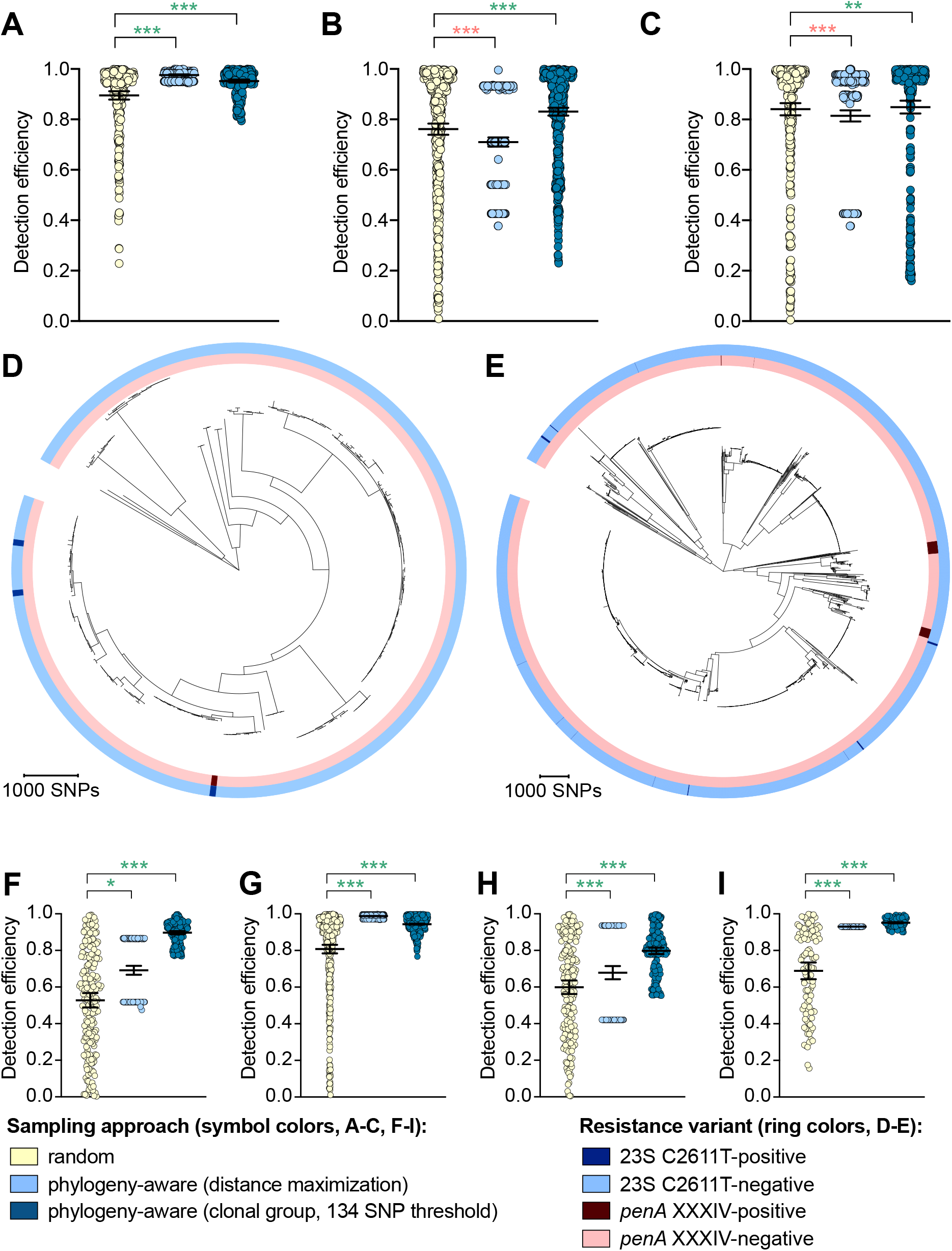
The impact of phylogeny-aware sampling on the detection efficiency of genetic resistance and diagnostic escape variants. Scatter dot plots showing the detection efficiency (with lines indicating the mean and 95% confidence intervals from 100 simulations) for resistance variants RplD G70D (**A**), 23S rRNA C2611T (**B**), and *penA* XXXIV (**C**) in datasets 1-5. Note that sampling simulations were not performed for RplD G70D in datasets 1 and 4 or for *penA* XXXIV in dataset 3 as prevalence of the variants in these datasets was >10%. Maximum-likelihood phylogenies produced from pseudogenome alignments (with predicted regions of recombination removed) of isolates from dataset 4 (**D**) and dataset 2 (**E**). Presence or absence of the 23S rRNA C2611T mutation (in at least 2/4 alleles) and the mosaic *penA* XXXIV allele is indicated by colored rings. Scatter dot plots showing the detection efficiency (with lines indicating the mean and 95% confidence intervals from 100 simulations) for diagnostic-associated variants 16S rRNA C1209A (**F**), *N. meningitidis-like porA* (**G**), *cppB* deletion (**H**), and DR-9A G168A (**I**) in all datasets in which the variant was present. Dot colors in **A-C** and **F-I** indicate the sampling approach, and asterisks indicate a significant difference (*P* < 0.05 by Mann-Whitney U test) in detection efficiency between the phylogeny-aware approach compared to random sampling (\**P* < 0.05, \*\**P* < 0.01, \*\*\**P* < 0.001; red asterisks indicate significantly lower detection efficiency of the phylogeny-aware approach compared to random sampling, and green asterisks indicate significantly higher detection efficiency of the phylogeny-aware approach compared to random sampling). n.s., not significant at α = 0.05.

The clonal group sampling approach prevents repeated sampling of very closely related isolates until all unique phylogenetic clusters have been sampled. Thus, for both rare variants that are clonally distributed and rare variants that are more randomly dispersed throughout the phylogeny (*e.g.*, *penA* XXXIV and 23S rRNA C2611T mutations, respectively, **Table 2**), this approach increases detection efficiency in cases where 1) there is substantial clonality among isolates and 2) a substantial proportion of variant-positive isolates do not occur in clonal lineages dominated by variant-negative isolates (**Fig. 2E**). In such datasets, effectively collapsing large variant-negative lineages into a single representative increases the effective prevalence of the variants and thus the detection efficiency of the clonal group approach compared to random sampling. The clonal group sampling approach significantly decreased detection efficiency in only one instance (*i.e.*, the 23S rRNA C2611T variant in dataset 4, **Table S5**), where all isolates with the variant appeared in large clonal lineages of predominately variant-negative isolates (**Fig. 2D**).

In cases where the clonal group sampling approach did not perform better than random sampling, adjusting the threshold for clonal grouping and/or a marginal increase in the prevalence of variant-positive isolates could elevate the relative performance of this targeted approach. We chose 134 SNPs as an example threshold for clonal grouping, as it represents the lower 95% confidence interval of the mean of SNP distances between each CFX-R resistant and the closest susceptible isolate in datasets 1-5 (see Methods). In the case of the 23S rRNA C2611T variant in dataset 4, the average prevalence of the variant across clonal groups (*i.e.*, the total number of variant-positive isolates, counting each variant-positive isolate as [1 / [1 + the total number of additional isolates that are ≤ 134 SNPs of the isolate]], divided by the number of clonal groups) is 0.005, lower than the actual prevalence of 0.012. However, if the threshold for clonal grouping was lower in this instance (*e.g.*, 50 SNPs), the effective prevalence of the variants would be 0.020, greater than the actual prevalence of 0.012. Similarly, using the 134 SNP threshold, if one additional isolate that was > 134 SNPs from any other isolates in this dataset had the 23S rRNA C2611T mutation, the average prevalence of the variant across clonal groups would be 0.036, greater than the actual prevalence of 0.016, and thus the clonal group approach would outperform random sampling.

To further assess the performance of phylogeny-aware sampling in the context of rare genetic variants that may have emerged in response to diagnostic pressure, we simulated random and phylogeny-aware sampling to assess detection efficiency of an additional set of variants. Specifically, we assessed a panel of *N. gonorrhoeae* diagnostic escape variants: the 16S rRNA C1209A mutation, the *N. meningitidis-like porA*, and the *cppB* deletion, all of which have been previously associated with diagnostic failure ^7–10^ and were present in one or more of datasets 1-5 at low prevalence (**Table 3**). The G168A mutation in the primer binding region of DR-9A, the target of the COBAS 4800 CT/NG (Roche) diagnostic, has not previously been documented but was present in 0.1% of strains from dataset 2. All of the diagnostic-associated variants assessed appeared in divergent backgrounds and were thus detected more efficiently by phylogeny-aware sampling compared to random sampling (**Fig. 2F-I, Table S6**). Like the results from the simulations based on resistance variants, the distance maximization approach maximized detection efficiency for some of the diagnostic-associated variants, but superiority of this approach to random sampling was not consistent across all variants. However, the clonal group approach performed significantly better than random sampling for all diagnostic-associated variants across all datasets.

**Table 3.** Summary of the potential diagnostic escape variants assessed

| Variant | Diagnostic assay | Documented association with diagnostic failure | Prevalence in dataset |  |  |  |  |
| --- | --- | --- | --- | --- | --- | --- | --- |
|  |  |  | 1 | 2 | 3 | 4 | 5 |
| 16S rRNA C1209A (4 alleles) | Aptima GC Combo | Yes<sup>7</sup> | 0.11% | 0.09% | 0% | 0% | 0% |
| *N. meningitidis*-like *porA* | In-house<sup>39,40</sup> | Yes<sup>8,9</sup> | 0.11% | 0.05% | 0% | 0% | 0% |
| *cppB* deletion | In-house<sup>41,42</sup> | Yes<sup>10</sup> | 1.12% | 0.05% | 0.47% | 0% | 7.29% |
| DR-9A G168A | Roche COBAS 4800 CT/NG | No | 0% | 0.09% | 0% | 0% | 0% |

The relative performance of the clonal group sampling approach compared to random sampling was generally consistent across multiple thresholds based on pseudogenomes (*i.e.*, ≤ 134 SNPs, ≤ 422 SNPs, and fastBAPS groups); relative performance of clonal group sampling using MLSTs, however, was less consistent and was significantly worse than random sampling for several variants (**Fig. S2, Tables S5-S6**). Together, these results suggest that preferentially sampling isolates that, based on whole genome sequencing (WGS), are phylogenetically divergent from those that have previously been sampled may increase detection efficiency of novel resistance variants.

### Targeted sampling based on genetic markers

Multiple drug resistance is more common in pathogenic bacteria than one would expect from the product of frequencies of resistance to individual drugs ^43,44^. This suggests that novel resistance mechanisms might be more likely to arise and spread in bacterial strains that are already resistant to other drugs, a phenomenon that has been documented in *N. gonorrhoeae* ^45^. It may therefore be fruitful to look for novel resistance variants for one drug in genetic backgrounds that are resistant to other drugs. It may be similarly effective to sample preferentially isolates with genetic markers that have been associated with a range of resistance mechanisms (*e.g.*, through epistatic interactions with other genetic variants) within and/or across different antibiotics when screening for a novel resistance variant. For example, as ciprofloxacin was the recommended first-line therapy for uncomplicated gonorrhea through 2005 in the United Kingdom ^46^, 2007 in the United States ^47^, and more recent years in other countries ^48–50^, we investigated whether resistance to ESCs is significantly more likely to occur in the background of genotypic ciprofloxacin resistance (*i.e.*, in strains with the GyrA S91F mutation). Similarly, as mutations at positions 120 and/or 121 in PorB, the major outer membrane protein in gonococci, have been associated with resistance to a range of drugs from multiple classes ^51^, we investigated whether resistance to ESCs is significantly more likely to occur in strains with PorB 120 and/or 121 mutations. Isolates with CRO-RS and CFX-R were significantly more likely to have the GyrA S91F mutation and the PorB G120 and/or A121 mutations than the wild-type GyrA S91 or wild-type PorB G120/A121 (*P* < 0.001, Fisher’s exact test, **Fig. 3A-B**). Further, both GyrA S91F and PorB G120 and/or A121 mutations occurred across a range of ESC resistance locus haplotypes (**Fig. 3C-D**). For all datasets with CRO-RS or CFX-R isolates, detection efficiency of both variants was significantly increased by only sampling isolates with the GyrA S91F mutation or the PorB G120 and/or A121 mutations (**Fig. 3E-F, Table S7**). Together, these results suggest that preferential sampling of isolates with certain genetic markers, including markers of resistance to previous first-line antibiotics, may increase the detection efficiency of novel resistance variants.

**Figure 3.**
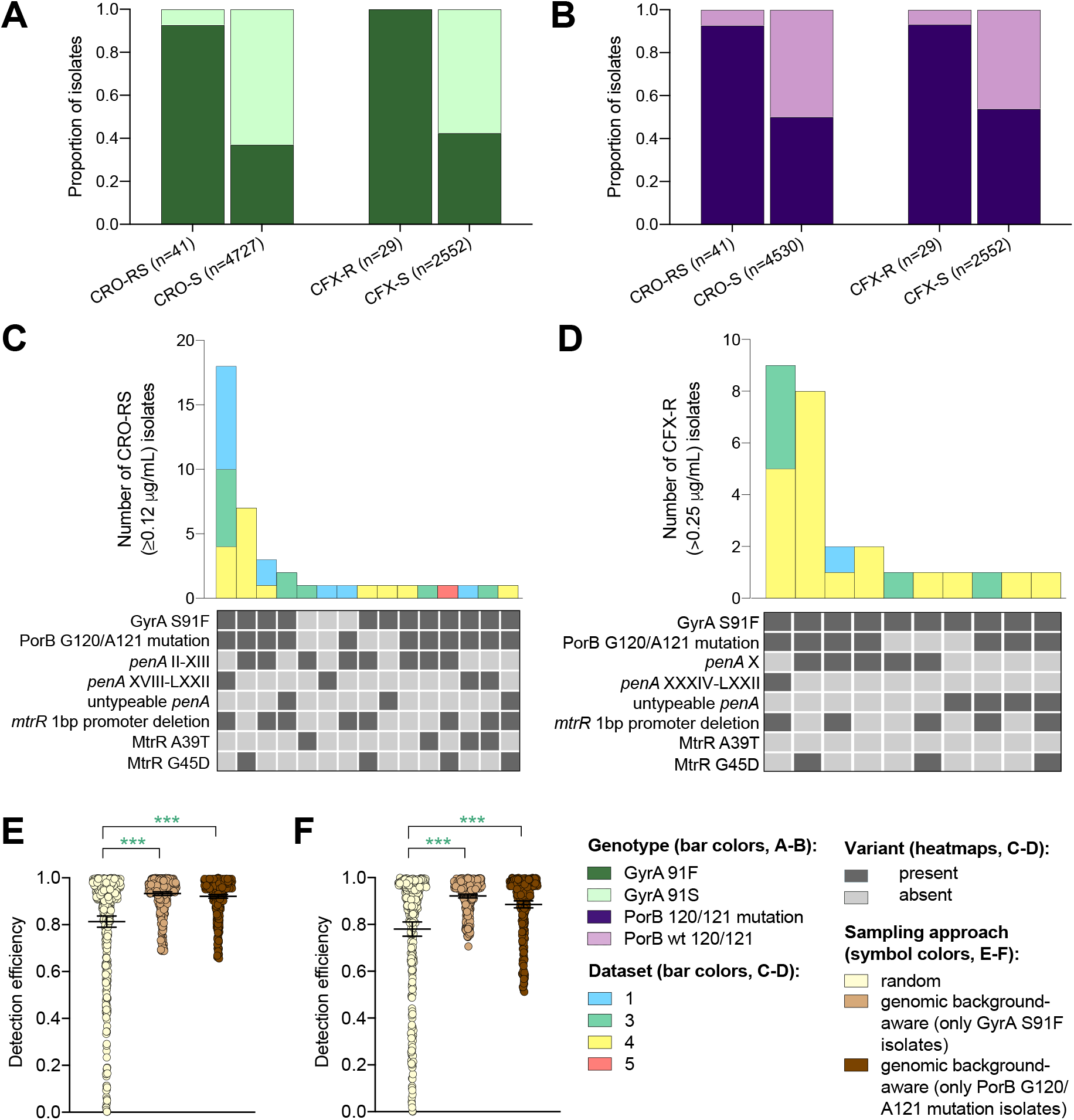
The impact of genomic background-aware sampling on the detection efficiency of phenotypic resistance variants. Bar charts showing the proportions of ceftriaxone reduced susceptibility (CRO-RS) isolates, ceftriaxone susceptible (CRO-S) isolates, cefixime resistant (CFX-R) isolates, and cefixime susceptible (CFX-S) isolates with GyrA S91F and GyrA S91 wild-type alleles (**A**) and with PorB G120 and/or A121 mutations and PorB G120 and A121 wild-type alleles (**B**) across datasets 1-5. Bar charts showing the number of (**C**) CRO-RS and (**D**) CFX-R isolates with each haplotype, along with heatmaps showing the presence or absence of the GyrA S19F mutation, the PorB G120 and/or A121 mutations, and other alleles at loci previously associated with extended spectrum cephalosporin resistance. Bar colors in (**C**) and (**D**) indicate the dataset from which the isolates were derived. Scatter dot plots showing the detection efficiency (with lines indicating the mean and 95% confidence intervals from 100 simulations) for CRO-RS (**E**) and CFX-R (**F**) in all datasets in which the variant was present. Dot colors in **E-F** indicate the sampling approach, and asterisks indicate a significant difference (*P* < 0.05 by Mann-Whitney U test) in detection efficiency between the phylogeny-aware approach compared to random sampling (\**P* < 0.05, \*\**P* < 0.01, \*\*\**P* < 0.001; green asterisks indicate significantly higher detection efficiency of the genomic background-aware approach compared to random sampling).

## Discussion

With sequencing becoming more integral to routine pathogen surveillance and diagnostics, it is important to ensure that models mapping genotypic information to expected pathogen phenotype and/or clinical outcome are comprehensive and current ^52^. In the case of genotype-based diagnostics, sustained phenotypic surveillance is crucial for identifying resistance variants that have recently emerged and/or increased in prevalence from previously undetected levels. While effective incorporation of patient metadata into surveillance strategies may be challenging, availability and incorporation of information on pathogen characteristics (*e.g.*, pathogen genomic data) into surveillance programs may ultimately decrease the cost of surveillance to maintain the sensitivity of these diagnostic tools.

Collection of patient metadata, including demographic and geographic information, is crucial to understanding the epidemiology of drug resistance. However, it may be difficult to obtain data on the relevant patient features, and the predictive power of such features may rapidly decay because of patient mobility and interactions ^53^. While availability of patient metadata varied across the datasets assessed, our results suggest that while incorporation of patient metadata into sampling strategies may increase detection efficiency for some novel resistance variants, it may be difficult to generalize for all potential novel resistance variants. It is possible that targeted sampling based on patient characteristics may be more reliable in the context of pathogens, antibiotic, and/or patient characteristics not assessed here.

Incorporation of WGS into routine pathogen surveillance by public health agencies ^54,55^ may facilitate use of genomic information in phenotypic sampling strategies, particularly with emerging metagenomic approaches that do not require bacterial culture ^56^. Our results show that phylogeny-aware sampling, particularly the clonal group approach, which reduces the amount of repeated sampling of closely related isolates, significantly improved detection efficiency over random sampling for multiple resistance and diagnostic-associated variants. Further, identification of and preferential sampling of isolates with genetic markers that are consistently predictive of resistance across a range of mechanisms, including those associated with resistance to other drugs, may supplement phylogeny-aware sampling to further optimize detection efficiency of novel variants. However, the utility of sampling based on genetic markers of other resistance mechanisms will likely vary substantially across different drugs and be influenced by future treatment guidelines.

While the clonal group sampling approach increased detection efficiency for the resistance and diagnostic escape variants assessed here, it may be difficult to determine the most effective and reliable metric or threshold for clonal grouping, especially as this is likely to vary across different clinical populations, antibiotics, and bacterial species. Detection efficiency was generally consistent across the two SNP thresholds and fastBAPS groupings based on WGS. However, performance of the clonal group approach using MLSTs was inconsistent and, in some instances, worse than random sampling, likely due to the shortcomings of MLST compared to WGS-based approaches in distinguishing between AMR variant-positive clades and more distantly-related variant-negative clades in species such as *N. gonorrhoeae* ^34^. This suggests that this approach is sensitive to similarity thresholds and that a low SNP threshold based on WGS assemblies may be the most appropriate approach, particularly in a population where there is expected to be substantial clonality among isolates and thus, even with a low threshold, detection efficiency will be improved by the clonal group approach. More broadly, surveillance incorporating WGS rather than MLST loci alone may further promote NAAT sustainability by enabling screening for variants with previously undetected mutations in target loci, such as the *N. gonorrhoeae* DR-9A G168A variants, that may be associated with diagnostic escape.

We have assessed these targeted sampling approaches in detection of multiple resistance variants across a range of populations, but these represent only a fraction of resistance mechanisms in a single species. These findings may extend to other antibiotics and bacterial species. For example, given the high degree of clonality among *M. tuberculosis* isolates and the significant variation in prevalence of drug resistance and resistance-conferring genotypes across clonal groups ^57,58^, the clonal group sampling approach may similarly improve detection efficiency of novel resistance variants in *M. tuberculosis*. For species in which drug resistance is primarily acquired through gene acquisition, it is unclear if phylogeny-aware sampling based on the core genome will improve detection efficiency of novel variants. K-mer distances ^59,60^ may provide a more practical alternative generalizable to more resistance mechanisms associated with gene acquisition. Further, the requirement of confirmatory phenotyping to identify novel resistance may not extend to pathogens that are expected to be associated with reliably-identifiable treatment failures, as for these pathogens, identification of treatment failure likely represents the most efficient method of novel resistance variant detection ^61^. However, for other pathogens, such as *N. gonorrhoeae* ^62^, treatment failures may go undetected for reasons including partial abatement of symptoms or long treatment regimens. Ultimately, as genotype-based diagnostics for antibiotic resistance become available for more species, it will be important to assess the efficiencies of these approaches across pathogens with different clinical, epidemiological, and evolutionary paradigms.

Since we lack the datasets to assess targeted sampling of variants from the time they first emerged in a population, any associations we observed between the variants and patient or pathogen features do not necessarily reflect those around the time of emergence. Thus, more longitudinal epidemiological and genomic studies, particularly after the implementation of genotype-based diagnostics, are necessary to better characterize patterns of novel resistance emergence and inform targeted surveillance approaches.

The phylogeny-aware sampling approaches presented here are based on the assumption that genomic data will be available for the pool of potential isolates from incident cases that may undergo confirmatory phenotyping. However, using information on isolate features to increase surveillance efficiency may be feasible even in the absence of mass prospective sequencing. For example, under the general assumption that novel resistance variants are more likely to appear in underrepresented lineages, phylogeny-aware surveillance could be paired with a diagnostic approach such as genomic neighbor typing ^56^, where any isolates with either susceptible or low confidence calls that appear to be divergent from the genomes in the reference database would be prioritized for confirmatory phenotyping. Similarly, a diagnostic that predicts AMR phenotypes through a combination of transcriptomic and genomic typing ^63^ may facilitate targeted surveillance by identifying isolates with ambiguous predictions (*e.g.*, isolates with transcriptional signatures of resistance that lack known genomic markers of resistance) that could be prioritized for confirmatory phenotyping.

Advances in diagnostics, extensive sequencing of clinical isolates, and large collections of clinical and pathogen data together provide new opportunities for integrating data streams and optimizing surveillance efforts. As marker-based point-of-care AMR diagnostics are developed and implemented, optimization of surveillance systems will require assessments like those modeled here of species-, drug-, and population-specific factors that may affect the emergence and distribution of diagnostic escape resistance variants, as well as how the diagnostic itself may complement surveillance efforts.

## Methods

### Dataset preparation and phylogenetic reconstruction

See **Table 1** for details of the *N. gonorrhoeae* datasets and **Tables 2** and **3** for the variants assessed. Raw sequencing data were downloaded from the NCBI Sequence Read Archive. Genomes were assembled using SPAdes v3.13 ^64^ with default parameters and the careful option to minimize the number of mismatches. Assembly quality was assessed using QUAST v4.3 ^65^, and contigs <500 bp in length and/or with <10x average coverage were removed. Isolate reference-based pseudogenomes were constructed by mapping raw reads to the NCCP11945 reference genome (RefSeq accession number NC_011035.1) using BWA-MEM v7.12 ^66^, the Picard toolkit v2.8 (http://broadinstitute.github.io/picard) to identify duplicate reads, and Pilon v1.22 ^67^ to determine the base call for each site, with a minimum depth of 10 and a minimum base quality of 20.

Loci in **Tables 2** and **3** were extracted from the genome assemblies using blastn
^68^ followed by MUSCLE alignment using default parameters ^69^ to assess the presence or absence of the resistance variants. Presence or absence of mutations in the multi-copy 16S and 23S rRNA genes and the repetitive DR-9A and DR-9B regions ^70^ was assessed using BWA-MEM, the Picard toolkit, and Pilon, as above, to map raw reads to a single 16S rRNA allele, a single 23S rRNA allele, a single DR-9A region, and a single DR-9B region from the NCCP11945 reference isolate and determine the mapping quality-weighted percentage of each nucleotide at the site of interest. Isolate metadata and resistance variant profiles are given in **Table S1.**

Gubbins v2.3.4 ^71^ was used with default parameters to identify and mask recombinant regions from the pseudogenomes and build maximum likelihood phylogenies from the non-recombinant pseudogenome alignments for each dataset through RAxML v8.2.12 ^72^. Pairwise phylogenetic distances were calculated after removal of predicted recombinant regions using the ape package in R. Phylogenetic distributions

73 of genetic resistance variants were assessed by estimating the phylogenetic D statistic ^73^ using the caper package in R. Bayesian analysis of population structure was performed on the pseudogenome alignments for each dataset using fastBAPS ^74^. Multilocus sequence types (MLSTs) were assigned using the PubMLST database (https://pubmlst.org/neisseria/).

### Sampling approaches

For each sampling approach/dataset/variant combination, 100 simulations were carried out with isolate sampling continuing until variant detection. We defined ‘detection efficiency’ as 1 minus the fraction of isolates sampled prior to variant detection (excluding any samples for which the presence or absence of the variant could not be determined). Because the purpose of this study was to compare the rare variant detection efficiency between random sampling and targeted sampling approaches, we did not evaluate RplD G70D in datasets 1 and 4 or for the *penA* XXXIV allele in dataset 3, as the prevalence of these variants in these datasets was > 10%.

In demography-aware sampling (datasets 1 and 2), the first isolate was selected at random, and each successive isolate was randomly selected from alternating demographic groups (men vs. women and men who have sex with men [MSM] vs. men who have sex with women [MSW] or women who have sex with men [WSM]). For anatomical site (niche)-aware sampling (datasets 1 and 2), the first isolate was selected at random, and each successive isolate was randomly selected from alternating anatomical sites of isolate collection (*i.e.*, cervix, urethra, rectum, and pharynx). For geography-aware sampling (datasets 3 and 4), the first isolate was selected at random, and each successive isolate was randomly selected from alternating geographic regions (countries or prefectures). For geography- and distance-aware sampling (datasets 3 and 4), the first isolate was selected at random, and each successive isolate was selected randomly from the region (country or prefecture) with the largest product of geographic distances from previously sampled regions, only re-sampling from a given region after all regions had been sampled in that round. For travel history- and sex work-aware sampling (dataset 2), isolates were selected at random either limiting the pool to isolates from patients who had recently engaged in overseas sex or sex work, respectively ^33^.

For phylogeny-aware sampling (datasets 1-5), the first isolate was selected at random, and each successive isolate was either selected to maximize the product of phylogenetic distances from each of the previously sampled isolates (“distance maximization”) or selected randomly with the exception of ensuring even sampling across phylogenetic groups (“clonal group”; *i.e.*, isolates ≤ *N* SNPs from a previously sampled isolate that were excluded from future sampling until all “clonal groups” had been sampled). SNP cutoffs tested for the clonal group approach included 1) 134 SNPs, the lower 95% confidence interval of the mean SNP distance across datasets 1-5 between each isolate with phenotypic cefixime resistance (CFX-R), azithromycin resistance (AZMR), and/or ceftriaxone reduced susceptibility (CRO-RS, >0.25 μg/mL, >1 μg/mL, and ≥0.12 μg/mL, respectively) and the closest susceptible isolate, and 2) 422 SNPs, the lower 95% confidence interval of the mean SNP distance across datasets 1-5 between each isolate with the RplD G70D mutation, the 23S rRNA C2611T mutation, and/or the *penA* XXXIV allele and the closest isolate without the resistance variant. The clonal group sampling approach was further tested by alternating sampling across fastBAPS and MLST groups.

For genomic background-aware sampling, isolates were selected at random either limiting the pool to isolates with genotypic ciprofloxacin resistance (*i.e.*, the GyrA S91F mutation) or to isolates with a mutation at PorB G120 and/or PorB A121, which have been associated with a range of resistance pathways in multiple classes of antibiotics ^51^.

Genomic background-aware sampling was assessed in detection of CRO-RS (datasets 1 and 3-5; dataset 2 had no CRO-RS isolates) and CFX-R (datasets 1 and 3-4; datasets 2 and 5 had no CFX-RS isolates).

## References

1 McAdams, D., Waldetoft, K. W., Tedijanto, C., Lipsitch, M. & Brown, S. P. Resistance diagnostics as a public health tool to combat antibiotic resistance: A model-based evaluation. PLoS Biol 17, e3000250, doi:https://doi.org/10.1371/journal.pbio.3000250 (2019).

2 Fingerhuth, S. M., Low, N., Bonhoeffer, S. & Althaus, C. L. Detection of antibiotic resistance is essential for gonorrhoea point-of-care testing: a mathematical modelling study. BMC Med 15, 142, doi:10.1186/s12916-017-0881-x (2017).

3 Tuite, A. R. et al. Impact of Rapid Susceptibility Testing and Antibiotic Selection Strategy on the Emergence and Spread of Antibiotic Resistance in Gonorrhea. J Infect Dis 216, 1141–1149, doi:10.1093/infdis/jix450 (2017).

4 Andre, E. et al. Novel rapid PCR for the detection of Ile491Phe rpoB mutation of Mycobacterium tuberculosis, a rifampicin-resistance-conferring mutation undetected by commercial assays. Clin Microbiol Infect 23, 267 e265–267 e267, doi:10.1016/j.cmi.2016.12.009 (2017).

5 Berhane, A. et al. Major Threat to Malaria Control Programs by Plasmodium falciparum Lacking Histidine-Rich Protein 2, Eritrea. Emerg Infect Dis 24, 462-470, doi:10.3201/eid2403.171723 (2018).

6 Herrmann, B. et al. Emergence and spread of Chlamydia trachomatis variant, Sweden. Emerg Infect Dis 14, 1462–1465, doi:10.3201/eid1409.080153 (2008).

7 Guglielmino, C. J. D., Appleton, S., Vohra, R. & Jennison, A. V. Identification of an unusual 16S rRNA mutation in Neisseria gonorrhoeae. J Clin Microbiol, doi:10.1128/JCM.01337-19 (2019).

8 Whiley, D. M. et al. False-negative results using Neisseria gonorrhoeae porA pseudogene PCR - a clinical gonococcal isolate with an N. meningitidis porA sequence, Australia, March 2011. Euro Surveill 16 (2011).

9 Golparian, D., Johansson, E. & Unemo, M. Clinical Neisseria gonorrhoeae isolate with a N. meningitidis porA gene and no prolyliminopeptidase activity, Sweden, 2011: danger of false-negative genetic and culture diagnostic results. Euro Surveill 17 (2012).

10 Bruisten, S. M. et al. Multicenter validation of the cppB gene as a PCR target for detection of Neisseria gonorrhoeae. J Clin Microbiol 42, 4332–4334, doi:10.1128/JCM.42.9.4332-4334.2004 (2004).

11 Lee, G. H., Pang, S. & Coombs, G. W. Misidentification of Staphylococcus aureus by the Cepheid Xpert MRSA/SA BC Assay Due to Deletions in the spa Gene. Journal of clinical microbiology 56, doi:10.1128/JCM.00530-18 (2018).

12 Marks, M. et al. Diagnostics for Yaws Eradication: Insights From Direct Next-Generation Sequencing of Cutaneous Strains of Treponema pallidum. Clin Infect Dis 66, 818–824, doi:10.1093/cid/cix892 (2018).

13 Smid, J. H., Althaus, C. L., Low, N., Unemo, M. & Herrmann, B. Rise and fall of the new variant of Chlamydia trachomatis in Sweden: mathematical modelling study. Sex Transm Infect, doi:10.1136/sextrans-2019-054057 (2019).

14 Hicks, A. L., Kissler, S. M., Lipsitch, M. & Grad, Y. H. Surveillance to maintain the sensitivity of genotype-based antibiotic resistance diagnostics. PLoS Biol 17, e3000547, doi:10.1371/journal.pbio.3000547 (2019).

15 Rempel, O. R. & Laupland, K. B. Surveillance for antimicrobial resistant organisms: potential sources and magnitude of bias. Epidemiol Infect 137, 16651673, doi:10.1017/S0950268809990100 (2009).

16 Unemo, M. et al. World Health Organization Global Gonococcal Antimicrobial Surveillance Program (WHO GASP): review of new data and evidence to inform international collaborative actions and research efforts. Sex Health, doi:10.1071/SH19023 (2019).

17 Hutinel, M. et al. Population-level surveillance of antibiotic resistance in Escherichia coli through sewage analysis. Euro Surveill 24, doi:10.2807/1560-7917.ES.2019.24.37.1800497 (2019).

18 Van Goethem, N. et al. Status and potential of bacterial genomics for public health practice: a scoping review. Implement Sci 14, 79, doi:10.1186/s13012-019-0930-2 (2019).

19 Lewis, D. A. The role of core groups in the emergence and dissemination of antimicrobial-resistant N gonorrhoeae. Sex Transm Infect 89 Suppl 4, iv47–51, doi:10.1136/sextrans-2013-051020 (2013).

20 Collignon, P., Beggs, J. J., Walsh, T. R., Gandra, S. & Laxminarayan, R. Anthropological and socioeconomic factors contributing to global antimicrobial resistance: a univariate and multivariable analysis. Lancet Planet Health 2, e398–e405, doi:10.1016/S2542-5196(18)30186-4 (2018).

21 Frost, I., Van Boeckel, T. P., Pires, J., Craig, J. & Laxminarayan, R. Global Geographic Trends in Antimicrobial Resistance: The Role of International Travel. J Travel Med, doi:10.1093/jtm/taz036 (2019).

22 Hernando Rovirola, C. et al. Antimicrobial resistance in Neisseria gonorrhoeae isolates from foreign-born population in the European Gonococcal Antimicrobial Surveillance Programme. Sex Transm Infect, doi:10.1136/sextrans-2018-053912 (2020).

23 Kirkcaldy, R. D., Weston, E., Segurado, A. C. & Hughes, G. Epidemiology of gonorrhoea: a global perspective. Sex Health, doi:10.1071/SH19061 (2019).

24 Perrin, L., Kaiser, L. & Yerly, S. Travel and the spread of HIV-1 genetic variants. Lancet Infect Dis 3, 22–27 (2003).

25 Pham Thanh, D. et al. A novel ciprofloxacin-resistant subclade of H58 Salmonella Typhi is associated with fluoroquinolone treatment failure. Elife 5, e14003, doi:10.7554/eLife.14003 (2016).

26 Kingsley, R. A. et al. Epidemic multiple drug resistant Salmonella Typhimurium causing invasive disease in sub-Saharan Africa have a distinct genotype. Genome Res 19, 2279–2287, doi:10.1101/gr.091017.109 (2009).

27 Mac Aogain, M., Rogers, T. R. & Crowley, B. Identification of emergent bla CMY-2-carrying Proteus mirabilis lineages by whole-genome sequencing. New Microbes New Infect 9, 58–62, doi:10.1016/j.nmni.2015.11.012 (2016).

28 Borrell, S. & Gagneux, S. Strain diversity, epistasis and the evolution of drug resistance in Mycobacterium tuberculosis. Clin Microbiol Infect 17, 815–820, doi:10.1111/j.1469-0691.2011.03556.x (2011).

29 Gould, I. M. & MacKenzie, F. M. Antibiotic exposure as a risk factor for emergence of resistance: the influence of concentration. J Appl Microbiol 92 Suppl, 78S–84S (2002).

30 Hook, E. W., 3rd & Kirkcaldy, R. D. A Brief History of Evolving Diagnostics and Therapy for Gonorrhea: Lessons Learned. Clin Infect Dis 67, 1294–1299, doi:10.1093/cid/ciy271 (2018).

31 Yahara, K. et al. Genomic surveillance of Neisseria gonorrhoeae to investigate the distribution and evolution of antimicrobial-resistance determinants and lineages. Microb Genom 4, doi:10.1099/mgen.0.000205 (2018).

32 Wiesner, P. J., Tronca, E., Bonin, P., Pedersen, A. H. & Holmes, K. K. Clinical spectrum of pharyngeal gonococcal infection. N Engl J Med 288, 181–185, doi:10.1056/NEJM197301252880404 (1973).

33 Williamson, D. et al. Bridging of Neisseria Gonorrhoeae Across Diverse Sexual Networks in the HIV Pre-Exposure Prophylaxis (PrEP) Era: A Clinical and Molecular Epidemiological Study. Nature Communications 10, 3988 (2019).

34 Harris, S. R. et al. Public health surveillance of multidrug-resistant clones of Neisseria gonorrhoeae in Europe: a genomic survey. Lancet Infect Dis 18, 758768, doi:10.1016/S1473-3099(18)30225-1 (2018).

35 Lee, R. S. et al. Genomic epidemiology and antimicrobial resistance of Neisseria gonorrhoeae in New Zealand. J Antimicrob Chemother 73, 353–364, doi:10.1093/jac/dkx405 (2018).

36 Grad, Y. H. et al. Genomic Epidemiology of Gonococcal Resistance to Extended-Spectrum Cephalosporins, Macrolides, and Fluoroquinolones in the United States, 2000-2013. J Infect Dis 214, 1579–1587, doi:10.1093/infdis/jiw420 (2016).

37 Ng, L. K., Martin, I., Liu, G. & Bryden, L. Mutation in 23S rRNA associated with macrolide resistance in Neisseria gonorrhoeae. Antimicrob Agents Chemother 46, 3020–3025 (2002).

38 Grad, Y. H. et al. Genomic epidemiology of Neisseria gonorrhoeae with reduced susceptibility to cefixime in the USA: a retrospective observational study. Lancet Infect Dis 14, 220–226, doi:10.1016/S1473-3099(13)70693-5 (2014).

39 Whiley, D. M. et al. A new confirmatory Neisseria gonorrhoeae real-time PCR assay targeting the porA pseudogene. Eur J Clin Microbiol Infect Dis 23, 705–710, doi:10.1007/s10096-004-1170-0 (2004).

40 Whiley, D. M. et al. A real-time PCR assay for the detection of Neisseria gonorrhoeae in genital and extragenital specimens. Diagn Microbiol Infect Dis 52, 1–5, doi:10.1016/j.diagmicrobio.2004.12.011 (2005).

41 Diemert, D. J., Libman, M. D. & Lebel, P. Confirmation by 16S rRNA PCR of the COBAS AMPLICOR CT/NG test for diagnosis of Neisseria gonorrhoeae infection in a low-prevalence population. J Clin Microbiol 40, 4056–4059, doi:10.1128/jcm.40.11.4056-4059.2002 (2002).

42 Van Dyck, E., Smet, H., Van Damme, L. & Laga, M. Evaluation of the Roche Neisseria gonorrhoeae 16S rRNA PCR for confirmation of AMPLICOR PCR-positive samples and comparison of its diagnostic performance according to storage conditions and preparation of endocervical specimens. J Clin Microbiol 39, 2280–2282, doi:10.1128/JCM.39.6.2280-2282.2001 (2001).

43 Chang, H. H. et al. Origin and proliferation of multiple-drug resistance in bacterial pathogens. Microbiol Mol Biol Rev 79, 101–116, doi:10.1128/MMBR.00039-14 (2015).

44 Lehtinen, S., Blanquart, F., Lipsitch, M., Fraser, C. & with the Maela Pneumococcal, C. On the evolutionary ecology of multidrug resistance in bacteria. PLoS Pathog 15, e1007763, doi:10.1371/journal.ppat.1007763 (2019).

45 Goldstein, E. et al. Factors related to increasing prevalence of resistance to ciprofloxacin and other antimicrobial drugs in Neisseria gonorrhoeae, United States. Emerg Infect Dis 18, 1290–1297, doi:10.3201/eid1808.111202 (2012).

46 Whittles, L. K., White, P. J., Paul, J. & Didelot, X. Epidemiological Trends of Antibiotic Resistant Gonorrhoea in the United Kingdom. Antibiotics (Basel) 7, doi:10.3390/antibiotics7030060 (2018).

47 Centers for Disease, C. & Prevention. Update to CDC’s sexually transmitted diseases treatment guidelines, 2006: fluoroquinolones no longer recommended for treatment of gonococcal infections. MMWR Morb Mortal Wkly Rep 56, 332–336 (2007).

48 Hemarajata, P., Yang, S., Soge, O. O., Humphries, R. M. & Klausner, J. D. Performance and Verification of a Real-Time PCR Assay Targeting the gyrA Gene for Prediction of Ciprofloxacin Resistance in Neisseria gonorrhoeae. J Clin Microbiol 54, 805–808, doi:10.1128/JCM.03032-15 (2016).

49 Unemo, M. & Dillon, J. A. Mitigating the emergence and spread of multidrug- and extensively drug-resistant gonorrhea: is there sufficient support in resource-poor settings in Africa? Sex Transm Dis 41, 238–239, doi:10.1097/OLQ.0000000000000117 (2014).

50 Bazzo, M. L. et al. First nationwide antimicrobial susceptibility surveillance for Neisseria gonorrhoeae in Brazil, 2015-16. J Antimicrob Chemother 73, 1854-1861, doi:10.1093/jac/dky090 (2018).

51 Mortimer, T. D. & Grad, Y. H. Applications of genomics to slow the spread of multidrug-resistant Neisseria gonorrhoeae. Ann N Y Acad Sci 1435, 93–109, doi:10.1111/nyas.13871 (2019).

52 Dona, V., Low, N., Golparian, D. & Unemo, M. Recent advances in the development and use of molecular tests to predict antimicrobial resistance in Neisseria gonorrhoeae. Expert Rev Mol Diagn 17, 845–859, doi:10.1080/14737159.2017.1360137 (2017).

53 Goldstein, E., Pitzer, V. E., O’Hagan, J. J. & Lipsitch, M. Temporally Varying Relative Risks for Infectious Diseases: Implications for Infectious Disease Control. Epidemiology 28, 136–144, doi:10.1097/EDE.0000000000000571 (2017).

54 European Centre for Disease Prevention and Control. ECDC strategic framework for the integration of molecular and genomic typing into European surveillance and multi-country outbreak investigations. (2019).

55 Brown, E., Dessai, U., McGarry, S. & Gerner-Smidt, P. Use of Whole-Genome Sequencing for Food Safety and Public Health in the United States. Foodborne Pathog Dis 16, 441–450, doi:10.1089/fpd.2019.2662 (2019).

56 Břinda, K. et al. Rapid heuristic inference of antibiotic resistance and susceptibility by genomic neighbor typing. bioRxiv, doi:http://dx.doi.org/10.1101/403204 (2019).

57 Merker, M. et al. Evolutionary history and global spread of the Mycobacterium tuberculosis Beijing lineage. Nat Genet 47, 242–249, doi:10.1038/ng.3195 (2015).

58 Casali, N. et al. Evolution and transmission of drug-resistant tuberculosis in a Russian population. Nat Genet 46, 279–286, doi:10.1038/ng.2878 (2014).

59 Ondov, B. D. et al. Mash: fast genome and metagenome distance estimation using MinHash. Genome Biol 17, 132, doi:10.1186/s13059-016-0997-x (2016).

60 Lees, J. A. et al. Fast and flexible bacterial genomic epidemiology with PopPUNK. Genome Res 29, 304–316, doi:10.1101/gr.241455.118 (2019).

61 Berenger, B. M. et al. Genetic Characterization and Enhanced Surveillance of Ceftriaxone-Resistant Neisseria gonorrhoeae Strain, Alberta, Canada, 2018. Emerg Infect Dis 25, 1660–1667, doi:10.3201/eid2509.190407 (2019).

62 Eyre, D. W. et al. Detection in the United Kingdom of the Neisseria gonorrhoeae FC428 clone, with ceftriaxone resistance and intermediate resistance to azithromycin, October to December 2018. Euro Surveill 24, doi:10.2807/1560-7917.ES.2019.24.10.1900147 (2019).

63 Bhattacharyya, R. P. et al. Simultaneous detection of genotype and phenotype enables rapid and accurate antibiotic susceptibility determination. Nat Med 25, 1858–1864, doi:10.1038/s41591-019-0650-9 (2019).

64 Bankevich, A. et al. SPAdes: a new genome assembly algorithm and its applications to single-cell sequencing. J Comput Biol 19, 455–477, doi:10.1089/cmb.2012.0021 (2012).

65 Gurevich, A., Saveliev, V., Vyahhi, N. & Tesler, G. QUAST: quality assessment tool for genome assemblies. Bioinformatics 29, 1072–1075, doi:10.1093/bioinformatics/btt086 (2013).

66 Li, H. Aligning sequence reads, clone sequences and assembly contigs with BWA-MEM. arXiv e-prints, doi:arXiv:1303.3997 (2013).

67 Walker, B. J. et al. Pilon: an integrated tool for comprehensive microbial variant detection and genome assembly improvement. PLoS One 9, e112963, doi:10.1371/journal.pone.0112963 (2014).

68 Altschul, S. F., Gish, W., Miller, W., Myers, E. W. & Lipman, D. J. Basic local alignment search tool. J Mol Biol 215, 403–410, doi:10.1016/S0022-2836(05)80360-2 (1990).

69 Edgar, R. C. MUSCLE: multiple sequence alignment with high accuracy and high throughput. Nucleic Acids Res 32, 1792–1797, doi:10.1093/nar/gkh340 (2004).

70 Kawa, D., Lu, S.-D. & Dailey, P. Oligonucleotides, methods and kits for detecting Neisseria Gonorrhoeae. wO patent EP1697541B1 (2013).

71 Croucher, N. J. et al. Rapid phylogenetic analysis of large samples of recombinant bacterial whole genome sequences using Gubbins. Nucleic Acids Res 43, e15, doi:10.1093/nar/gku1196 (2015).

72 Sommer, M. O. A., Munck, C., Toft-Kehler, R. V. & Andersson, D. I. Prediction of antibiotic resistance: time for a new preclinical paradigm? Nat Rev Microbiol 15, 689–696, doi:10.1038/nrmicro.2017.75 (2017).

73 Fritz, S. A. & Purvis, A. Selectivity in mammalian extinction risk and threat types: a new measure of phylogenetic signal strength in binary traits. Conserv Biol 24, 1042–1051, doi:10.1111/j.1523-1739.2010.01455.x (2010).

74 Tonkin-Hill, G., Lees, J. A., Bentley, S. D., Frost, S. D. W. & Corander, J. Fast hierarchical Bayesian analysis of population structure. Nucleic Acids Res 47, 5539–5549, doi:10.1093/nar/gkz361 (2019).

