## Supplementary data for "Targeted surveillance strategies for efficient detection of novel antibiotic resistance variants"

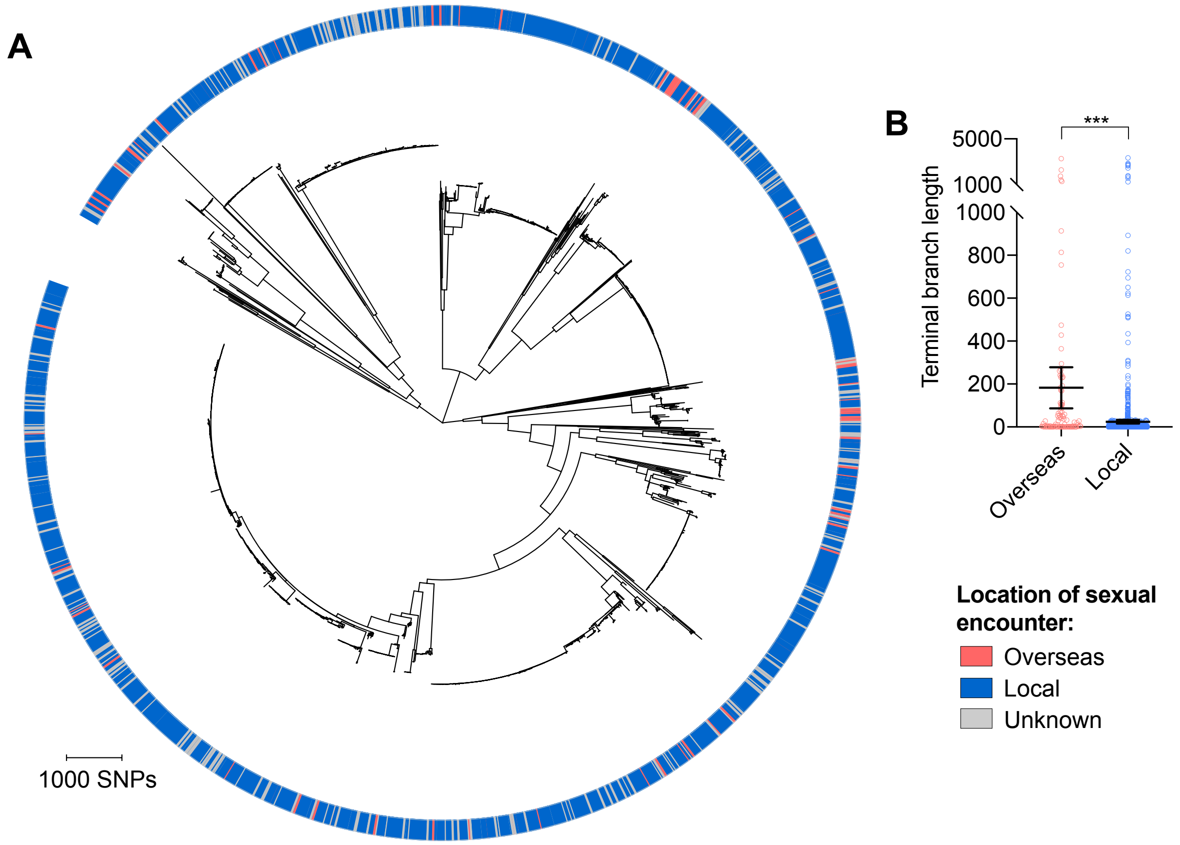


**Figure S1. Isolates from patients with travel-associated gonorrhea are associated with longer terminal branches compared to patients with locally-acquired gonorrhea.** Maximum-likelihood phylogeny produced from the pseudogenome alignment (with predicted regions of recombination removed) of isolates from dataset 2 (**A**). Patient travel history is indicated by the colored ring in **A**. Scatter dot plots showing the terminal branch lengths (with lines indicating the mean and 95% confidence intervals) associated with isolates from patients with travel-associated gonorrhea compared to patients with locally-acquired gonorrhea with asterisks indicating a significant difference ($P < 0.001$ by Mann-Whitney U test) in terminal branch lengths between the two groups (**B**).

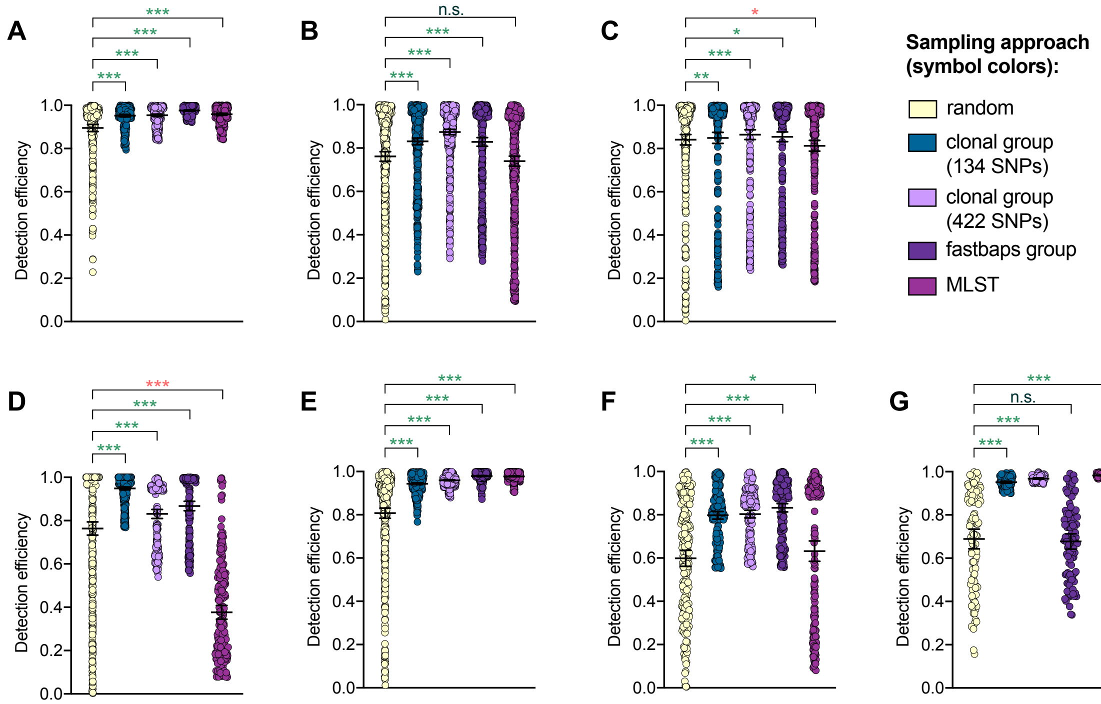


**Figure S2. Detection efficiency of clonal group sampling across different similarity thresholds.** Scatter dot plots showing the detection efficiency (with lines indicating the mean and 95% confidence intervals from 100 simulations) for resistance variants RplD G70D (**A**), 23S rRNA C2611T (**B**), and *penA* XXXIV (**C**) in datasets 1-5 and for diagnostic-associated variants 16S rRNA C1209A (**D**), *N. meningitidis*-like *porA* (**E**), *cppB* deletion (**F**), and DR-9A G168A (**G**) in all datasets in which the variant was present. Note that sampling simulations were not performed for RplD G70D in datasets 1 and 4 or for *penA* XXXIV in dataset 3 as prevalence of the variants in these datasets was >10%. Dot colors indicate the sampling approach, and asterisks indicate a significant difference ($P < 0.05$ by Mann-Whitney U test) in detection efficiency between the phylogeny-aware approach compared to random sampling (*$P < 0.05$, **$P < 0.01$, ***$P < 0.001$; red asterisks indicate significantly lower detection efficiency of the phylogeny-aware approach compared to random sampling, and green asterisks indicate significantly higher detection efficiency of the phylogeny-aware approach compared to random sampling). n.s., not significant at $\alpha = 0.05$.

**Table S2.** Demographic and geographic sampling biases in datasets 1-3.

| | | **Dataset 1 [Mortimer et al., 2020, *in preparation*]**[a] | **US (2011-2015)**[1-5] | **Dataset 2**[6a] | **Victoria, Australia (2017)** | **Dataset 3**[7] | **Euro-GASP countries (2013)**[8] |
|---|---|---|---|---|---|---|---|
| **Proportion of total gonorrhea cases that were from** | men | 0.958 | 0.603 | 0.872 | 0.81 | N/A | N/A |
| | MSM | 0.737 | 0.295 | 0.733 | 0.567 | N/A | N/A |
| | Austria | N/A | N/A | N/A | N/A | 0.051 | 0.023 |
| | Belgium | N/A | N/A | N/A | N/A | 0.052 | 0.021 |
| | Cyprus | N/A | N/A | N/A | N/A | 0.008 | 0.000 |
| | Denmark | N/A | N/A | N/A | N/A | 0.052 | 0.017 |
| | France | N/A | N/A | N/A | N/A | 0.054 | 0.028 |
| | Germany | N/A | N/A | N/A | N/A | 0.045 | N/A |
| | Greece | N/A | N/A | N/A | N/A | 0.046 | 0.004 |
| | Hungary | N/A | N/A | N/A | N/A | 0.046 | 0.031 |
| | Iceland | N/A | N/A | N/A | N/A | 0.005 | 0.000 |
| | Italy | N/A | N/A | N/A | N/A | 0.025 | 0.026 |
| | Latvia | N/A | N/A | N/A | N/A | 0.036 | 0.011 |
| | Malta | N/A | N/A | N/A | N/A | 0.019 | 0.001 |
| | Netherlands | N/A | N/A | N/A | N/A | 0.063 | 0.085 |
| | Norway | N/A | N/A | N/A | N/A | 0.052 | 0.010 |
| | Portugal | N/A | N/A | N/A | N/A | 0.102 | 0.002 |
| | Slovakia | N/A | N/A | N/A | N/A | 0.036 | 0.008 |
| | Slovenia | N/A | N/A | N/A | N/A | 0.051 | 0.001 |
| | Spain | N/A | N/A | N/A | N/A | 0.110 | 0.068 |
| | Sweden | N/A | N/A | N/A | N/A | 0.047 | 0.002 |
| | UK | N/A | N/A | N/A | N/A | 0.101 | 0.661 |

MSM, men who have sex with men

[a]Proportion of patients with identified gender or sexual behavior that identified as men or MSM

**Table S3.** Detection efficiency of random, demography-, niche-, and geography-aware sampling approaches for resistance variants.

| Dataset | Variant | Sampling approach[a] | | | | | |
|---|---|---|---|---|---|---|---|
| | | Random[b] | Demography-aware (M vs. W) | Demography-aware (MSM vs. WSM/MSW) | Niche-aware | Geography- and distance-aware | Geography-aware |
| 1 | 23S C2611T (2-4 alleles) | 0.51 (0.46-0.57) | 0.52 (0.46-0.57) | 0.45 (0.4-0.5) | 0.41 (0.36-0.47)* | N/A | N/A |
| | *penA* XXXIV | 0.98 (0.97-0.98) | 0.98 (0.98-0.98) | 0.98 (0.98-0.99) | 0.97 (0.97-0.98) | N/A | N/A |
| 2 | RplD G70D | 0.96 (0.96-0.97) | 0.96 (0.95-0.97) | 0.95 (0.94-0.96)* | 0.96 (0.95-0.97) | N/A | N/A |
| | 23S C2611T (2-4 alleles) | 0.97 (0.96-0.97) | 0.97 (0.96-0.97) | 0.96 (0.95-0.97) | 0.96 (0.95-0.97) | N/A | N/A |
| | *penA* XXXIV | 0.98 (0.97-0.98) | 0.96 (0.96-0.97)** | 0.96 (0.96-0.97)** | 0.97 (0.97-0.98) | N/A | N/A |
| 3 | RplD G70D | 0.96 (0.95-0.97) | N/A | N/A | N/A | 0.96 (0.96-0.97) | 0.96 (0.95-0.96) |
| | 23S C2611T (2-4 alleles) | 0.93 (0.91-0.94) | N/A | N/A | N/A | 0.88 (0.86-0.9)** | 0.88 (0.86-0.9)** |
| 4 | 23S C2611T (2-4 alleles) | 0.75 (0.71-0.79) | N/A | N/A | N/A | 0.67 (0.64-0.71)*** | 0.63 (0.6-0.66)*** |
| | *penA* XXXIV | 0.52 (0.46-0.58) | N/A | N/A | N/A | 0.42 (0.37-0.47)* | 0.43 (0.39-0.48)* |

[a]Mean detection efficiency with 95% confidence intervals. Asterisks indicate significantly different detection efficiency of targeted sampling approach compared to random sampling ($P < 0.05$ by Mann Whitney U Test). ***, $P < 0.001$; **, $P < 0.01$; *, $P < 0.05$.
[b]Mean detection efficiency with 95% confidence intervals achieved by random sampling from all isolates for which the presence of absence of the variant could be determined. Note, however, that for targeted sampling approaches based on each different patient characteristic, isolates with missing information for that patient characteristic were removed, and random sampling was simulated on the reduced dataset for comparison to the targeted sampling approach.

**Table S4.** Detection efficiency of random sampling, as well as preferential sampling of patients that had recently engaged in overseas sex or in sex work, for resistance variants in dataset 2.

| **Variant** | **Sampling approach**[a] | | |
|---|---|---|---|
| | **Random**[b] | **Only patients with recent overseas sex** | **Only sex workers** |
| 23S C2611T (2-4 alleles) | 0.96 (0.95-0.97) | 0[c] | N/A |
| *penA* XXXIV | 0.96 (0.96-0.97) | 0.98 (0.98-0.98)*** | N/A |
| RplD G70D | 0.96 (0.95-0.96) | 0.99 (0.98-0.99)*** | N/A |
| 23S C2611T (2-4 alleles) | 0.96 (0.95-0.96) | N/A | 0.98 (0.98-0.98)*** |
| *penA* XXXIV | 0.97 (0.96-0.97) | N/A | 0.98 (0.97-0.98) |
| RplD G70D | 0.96 (0.95-0.97) | N/A | 0.97 (0.97-0.98) |

[a]Mean detection efficiency with 95% confidence intervals. Asterisks indicate significantly different detection efficiency of targeted sampling approach compared to random sampling ($P < 0.05$ by Mann Whitney U Test). ***, $P < 0.001$; **, $P < 0.01$; *, $P < 0.05$.
[b]Mean detection efficiency with 95% confidence intervals achieved by random sampling from all isolates for which the presence of absence of the variant could be determined and the relevant patient metadata (*i.e.,* overseas vs. local sex or sex worker status) was available.
[c]No isolates with 23S C2611T mutations were from patients with recent overseas sex.

**Table S5.** Detection efficiency of random and phylogeny-aware sampling approaches for resistance variants.

| Dataset | Variant | Sampling approach[a] | | | | |
|---|---|---|---|---|---|---|
| | | **Random** | **Phylogeny-aware (distance maximization)** | **Phylogeny-aware (clonal group, 134 SNP threshold)** | **Phylogeny-aware (clonal group, 422 SNP threshold)** | **Phylogeny-aware (fastbaps groups)** |
| 1 | 23S C2611T (2-4 alleles) | 0.51 (0.46-0.57) | 0.54 (0.54-0.54) | 0.71 (0.68-0.74)*** | 0.87 (0.86-0.89)*** | 0.54 (0.5-0.58) |
| | *penA* XXXIV | 0.98 (0.97-0.98) | 0.9 (0.89-0.9)*** | 0.99 (0.98-0.99)* | 0.99 (0.98-0.99) | 0.97 (0.97-0.98) |
| 2 | RplD G70D | 0.96 (0.96-0.97) | 0.99 (0.99-0.99)*** | 0.99 (0.98-0.99)*** | 0.99 (0.99-0.99)*** | 0.99 (0.99-0.99)*** |
| | 23S C2611T (2-4 alleles) | 0.97 (0.96-0.97) | 0.93 (0.93-0.93) *** | 0.98 (0.98-0.99)*** | 0.99 (0.98-0.99)*** | 0.97 (0.96-0.97) |
| | *penA* XXXIV | 0.98 (0.97-0.98) | 0.98 (0.98-0.98)** | 0.99 (0.99-0.99)*** | 0.99 (0.99-0.99)*** | 0.99 (0.99-0.99)*** |
| 3 | RplD G70D | 0.96 (0.95-0.97) | 0.95 (0.95-0.95)*** | 0.97 (0.97-0.97) | 0.97 (0.96-0.97) | 0.98 (0.97-0.98)** |
| | 23S C2611T (2-4 alleles) | 0.93 (0.91-0.94) | 0.71 (0.71-0.71)*** | 0.91 (0.9-0.93) | 0.85 (0.84-0.87)*** | 0.98 (0.98-0.99)*** |
| 4 | 23S C2611T (2-4 alleles) | 0.75 (0.71-0.79) | 0.44 (0.42-0.45)*** | 0.64 (0.6-0.69)*** | 0.68 (0.64-0.72)* | 0.67 (0.63-0.71)*** |
| | *penA* XXXIV | 0.52 (0.46-0.58) | 0.43 (0.42-0.44)* | 0.48 (0.42-0.53) | 0.53 (0.48-0.58) | 0.49 (0.45-0.53) |
| 5 | RplD G70D | 0.76 (0.73-0.8) | 0.98 (0.98-0.98)*** | 0.9 (0.89-0.91)*** | 0.9 (0.89-0.91)*** | 0.96 (0.96-0.97)*** |
| | 23S C2611T (2-4 alleles) | 0.65 (0.6-0.7) | 0.93 (0.93-0.93)*** | 0.9 (0.89-0.91)*** | 0.91 (0.9-0.92)*** | 0.98 (0.97-0.98)*** |
| | *penA* XXXIV | 0.89 (0.87-0.91) | 0.95 (0.95-0.95)*** | 0.95 (0.94-0.95)*** | 0.9 (0.89-0.91)*** | 0.96 (0.96-0.97)*** |

[a]Mean detection efficiency with 95% confidence intervals. Asterisks indicate significantly different detection efficiency of targeted sampling approach compared to random. sampling ($P$ < 0.05 by Mann Whitney U Test). ***, $P$ < 0.001; **, $P$ < 0.01; *, $P$ < 0.05

**Table S6.** Detection efficiency of random and phylogeny-aware sampling approaches for variants associated with diagnostic escape.

| Dataset | Variant | Sampling approach[a] | | | | |
|---|---|---|---|---|---|---|
| | | **Random** | **Phylogeny-aware (distance maximization)** | **Phylogeny-aware (clonal group, 134 SNP threshold)** | **Phylogeny-aware (clonal group, 422 SNP threshold)** | **Phylogeny-aware (fastbaps groups)** |
| 1 | *N. meningitidis*-like porA | 0.52 (0.46-0.58) | 0.52 (0.52-0.52) | 0.86 (0.85-0.87)*** | 0.7 (0.68-0.73)*** | 0.74 (0.72-0.77)*** |
| | *cppB* deletion | 0.91 (0.9-0.93) | 0.99 (0.99-0.99)*** | 0.96 (0.95-0.96)*** | 0.97 (0.96-0.97)*** | 0.98 (0.98-0.99)*** |
| | 16S rRNA | 0.51 (0.45-0.56) | 0.42 (0.42-0.42)*** | 0.74 (0.71-0.77)*** | 0.71 (0.68-0.73)*** | 0.73 (0.71-0.76)*** |
| | DR-9A G168A | N/A | N/A | N/A | N/A | N/A |
| 2 | *N. meningitidis*-like porA | 0.53 (0.48-0.59) | 0.87 (0.87-0.87)*** | 0.93 (0.93-0.94)*** | 0.96 (0.95-0.96)*** | 0.99 (0.99-0.99)*** |
| | *cppB* deletion | 0.51 (0.45-0.56) | 1 (1-1)*** | 0.93 (0.93-0.94)*** | 0.96 (0.95-0.96)*** | 0.99 (0.99-0.99)*** |
| | 16S rRNA | 0.69 (0.64-0.73) | 0.94 (0.94-0.94)*** | 0.85 (0.84-0.87)*** | 0.9 (0.89-0.91)*** | 0.93 (0.93-0.94)*** |
| | DR-9A G168A | 0.69 (0.64-0.73) | 0.93 (0.93-0.93)*** | 0.95 (0.95-0.96)*** | 0.97 (0.97-0.97)*** | 0.68 (0.64-0.71)*** |
| 3 | *N. meningitidis*-like porA | N/A | N/A | N/A | N/A | N/A |
| | *cppB* deletion | 0.85 (0.82-0.87) | 0.99 (0.99-0.99)*** | 0.91 (0.9-0.92)*** | 0.94 (0.93-0.94)*** | 0.99 (0.98-0.99)*** |
| | 16S rRNA | N/A | N/A | N/A | N/A | N/A |
| | DR-9A G168A | N/A | N/A | N/A | N/A | N/A |
| 5 | *N. meningitidis*-like porA | N/A | N/A | N/A | N/A | N/A |
| | *cppB* deletion | 0.96 (0.96-0.97) | 0.97 (0.97-0.97) | 0.98 (0.97-0.98)** | 0.98 (0.97-0.98)* | 0.96 (0.95-0.96)** |
| | 16S rRNA | N/A | N/A | N/A | N/A | N/A |
| | DR-9A G168A | N/A | N/A | N/A | N/A | N/A |

[a]Mean detection efficiency with 95% confidence intervals. Asterisks indicate significantly different detection efficiency of targeted sampling approach compared to random sampling ($P < 0.05$ by Mann Whitney U Test). ***, $P < 0.001$; **, $P < 0.01$; *, $P < 0.05$

**Table S7.** Detection efficiency of random and genomic background-aware sampling approaches for resistance variants.

| Dataset | Variant | Sampling approach[a] | | |
|---|---|---|---|---|
| | | Random | Genomic background-aware (only GyrA S91F isolates) | Genomic background-aware (only PorB G120/A121 mutation isolates) |
| 1 | CRO-RS (≥0.12 μg/mL) | 0.91 (0.9-0.93) | 0.98 (0.97-0.98)*** | 0.96 (0.95-0.97)*** |
| | CFX-R (>0.25 μg/mL) | 0.51 (0.45-0.57) | 0.87 (0.85-0.88)*** | 0.77 (0.74-0.8)*** |
| 3 | CRO-RS (≥0.12 μg/mL) | 0.91 (0.89-0.93) | 0.95 (0.94-0.96)*** | 0.94 (0.93-0.95)* |
| | CFX-R (>0.25 μg/mL) | 0.88 (0.86-0.9) | 0.93 (0.92-0.94)*** | 0.92 (0.9-0.93)*** |
| 4 | CRO-RS (≥0.12 μg/mL) | 0.93 (0.92-0.94) | 0.95 (0.94-0.96)** | 0.96 (0.95-0.97)*** |
| | CFX-R (>0.25 μg/mL) | 0.96 (0.95-0.96) | 0.97 (0.96-0.97)* | 0.97 (0.97-0.98)* |
| 5 | CRO-RS (≥0.12 μg/mL) | 0.5 (0.45-0.56) | 0.85 (0.83-0.87)*** | 0.83 (0.81-0.85)*** |
| | CFX-R (>0.25 μg/mL) | N/A | N/A | N/A |

[a]Mean detection efficiency with 95% confidence intervals. Asterisks indicate significantly different detection efficiency of targeted sampling approach compared to random sampling ($P < 0.05$ by Mann Whitney U Test). ***, $P < 0.001$; **, $P < 0.01$; *, $P < 0.05$

### Supplementary References:


1 Centers for Disease Control and Prevention. Sexually Transmitted Disease Surveillance 2011. (U.S. Department of Health and Human Services, Atlanta, 2012).

2 Centers for Disease Control and Prevention. Sexually Transmitted Disease Surveillance 2012. (U.S. Department of Health and Human Services, Atlanta, 2013).

3 Centers for Disease Control and Prevention. Sexually Transmitted Disease Surveillance 2013. (U.S. Department of Health and Human Services, Atlanta, 2014).

4 Centers for Disease Control and Prevention. Sexually Transmitted Disease Surveillance 2014. (U.S. Department of Health and Human Services, Atlanta, 2015).

5 Centers for Disease Control and Prevention. Sexually Transmitted Disease Surveillance 2015. (U.S. Department of Health and Human Services, Atlanta, 2016).

6 Williamson, D. *et al.* Bridging of Neisseria Gonorrhoeae Across Diverse Sexual Networks in the HIV Pre-Exposure Prophylaxis (PrEP) Era: A Clinical and Molecular Epidemiological Study. *Nature Communications* **10**, 3988 (2019).

7 Harris, S. R. *et al.* Public health surveillance of multidrug-resistant clones of Neisseria gonorrhoeae in Europe: a genomic survey. *Lancet Infect Dis* **18**, 758-768, doi:10.1016/S1473-3099(18)30225-1 (2018).

8 European Centre for Disease Prevention and Control. Gonococcal antimicrobial susceptibility surveillance in Europe 2013. (ECDC, Stokholm, 2015).
